# Within-host evolution of drug tolerance in *Mycobacterium tuberculosis*

**DOI:** 10.1101/2025.07.29.667394

**Authors:** Valerie F. A. March, Kakha Mchedlishvili, Galo A. Goig, Nino Maghradze, Teona Avaliani, Rusudan Aspindzelashvili, Zaza Avaliani, Maia Kipiani, Nestani Tukvadze, Levan Jugheli, Selim Bouaouina, Anna Doetsch, Sevda Kalkan, Miriam Reinhardt, Sebastien Gagneux, Sonia Borrell

## Abstract

**Background:** *Mycobacterium tuberculosis* (Mtb) causes tuberculosis (TB) in humans. Poor treatment responses are a threat to global TB control, as such, understanding contributing factors to poor responses is important. We hypothesized that antibiotic tolerance could contribute to delayed culture conversion (recalcitrant TB), and resistance amplification in patients during TB treatment.

**Objectives:** To investigate the role of drug tolerance in delayed culture conversion and resistance amplification in TB patients.

**Methods:** We collected serial Mtb isolates from i) patients with drug-susceptible TB who remained culture positive for up to 6 years (i.e. recalcitrant TB), and ii) patients with multidrug-resistant TB (MDR-TB) where resistance amplified during treatment. We measured tolerance to rifampicin (RIF) in drug-susceptible TB strains and tolerance to moxifloxacin (MFX) in MDR-TB strains using a real-time time-kill assay.

**Results:** RIF tolerance evolved within-host, increasing up to and ~1.5-fold, however, there was no apparent contribution of RIF tolerance to delayed culture conversion. Tolerance to Mfx in MDR-TB patients appeared negatively associated with resistance amplification and consistently decreased over time in patients.

**Conclusion:** Our findings confirm that antibiotic tolerance evolves in Mtb within patients over time during treatment. However, there was no evidence that this tolerance influences treatment responses, calling for further investigation of contributors to adverse treatment responses and their mitigation.

## Introduction

With a reported ~1.2 million tuberculosis (TB) related deaths in 2023, *Mycobacterium tuberculosis* (Mtb) is the deadliest human infectious agent (1, 2). The standard of care to treat TB is combination antibiotic chemotherapy, which is highly efficacious for drug-susceptible TB (DS-TB)(1), however, drug resistance emerges and threatens TB control.

Multidrug-resistant TB (MDR-TB) characterized by resistance to frontline drugs rifampicin (RIF) and isoniazid, has historically been less successfully treated (1). Regimens to treat MDR-TB include second-line agents such as fluoroquinolones (3–5), and now incorporate novel anti-TB drugs (6–8), for which, alarmingly, strains harbouring resistance conferring mutations are already circulating (9). Resistance amplification, where drug-resistant strains gain further drug resistance, complicates patient treatment as there are limited efficacious second-line agents available.

Studies have shown that TB treatment outcomes are affected by sociodemographic factors such as sex and age (10–16) (10, 12, 16–19), alongside comorbidities such as HIV coinfection (11, 13, 15, 17, 18, 20, 21), or having a history of TB (10, 12–14, 16, 22). In addition, poor responses to treatment such as delayed culture conversion (failure to culture convert after 2-3 months) (10, 14) as well as drug resistance (23, 24) increase the risk of poor treatment outcomes in TB. Recently, we published a study using clinical and bacterial genomic data from Georgia, demonstrating bacterial determinants can contribute to treatment outcomes (25), thus setting a precedent to further explore the contribution of bacterial determinants to treatment outcomes.

In other diseases, drug tolerance, defined as the ability of bacteria to withstand extended durations in bactericidal antibiotics (26), has been demonstrated to contribute to chronic infection in patients (27) and the emergence of antibiotic resistance (28). The impact of tolerance in TB has been underexplored, despite being a phenotype that can develop within patients even under combination therapy (29).

With access to serial Mtb clinical isolates from historical TB patients at the National Centre for Tuberculosis and Lung Disease (NCTLD) in Tbilisi, Georgia, we investigated the hypothesis that antibiotic tolerance can contribute to poor treatment responses in TB. We examined the contribution of tolerance to RIF and fluoroquinolone moxifloxacin (Mfx) to delayed culture conversion (recalcitrant TB) and resistance amplification in strains from DS-TB and MDR-TB patients, respectively.

## Methods

### Ethical approval

The institutional Review Board of the NCTLD in Tbilisi, Georgia and the Ethics North- and Central Switzerland granted ethical approval for the use of the patient samples and data used in this study. The ethics committees waived the need for individual patient consent since only limited and anonymized clinical data were used.

### Strain selection

We explored a database of Georgian patient data and clinical strains spanning the period between January 2008 and December 2022 to identify TB cases with serial isolates exhibiting recalcitrance in DS-TB and resistance amplification in MDR TB alongside appropriate controls. We selected three DS-TB patients that were typically cured (Group A cases), and six patients which had positive sputum cultures for extended durations as examples of recalcitrant TB cases (Group B cases). Moreover, we selected MDR-TB patients whose drug resistance profile remained the same throughout treatment (Group C cases) and MDR-TB patients where there was evidence of resistance amplification (Group D cases). Our resulting cohort was comprised of 18 bacteriologically confirmed adult patients with active pulmonary TB. Strains from these patients were shipped to our BLSL3 facilities in Switzerland for processing and experiments.

### Preparation of antibiotic stocks

Antibiotic stocks were prepared by dissolving powdered stocks of rifampicin (RIF, Sigma-Aldrich) and moxifloxacin (Mfx, Sigma-Aldrich) in dimethyl sulfoxide (DMSO, Applichem) to a desired stock concentration. The same drug stocks were used for MIC measurements and time-kill assays.

### Measuring antibiotic tolerance

Tolerance to rifampicin (RIF) was measured for drug susceptible strains and tolerance to moxifloxacin (Mfx) was measured for MDR strains using our real-time time killing assay, as previously described(30). In short, 50 µl of thawed calibrated stocks of strains were inoculated into 96-well u-bottom deep-well plates (Thermo Fisher) containing 950 µl of 7H9 ADC with or without antibiotic at 400 X MIC in triplicate. Resulting cultures were mixed thoroughly before 100 µl was sampled for viability using the BacTiter-Glo assay in a 1:1 ratio. BacTiter-Glo plates were incubated for 30 – 35 minutes before measuring luminescence using the Tecan Spark Multimodal plate reader for susceptible strains and the Tecan Infinite Pro 200 for MDR strains. Killing was measured every 4 days for 14 days, with a final measurement at 21 days post-inoculation. For Mfx experiments, a measurement after two days was included.

After controlling for background luminescence by subtracting signal from culture-free negative controls, killing was assessed by calculating the fraction of survival for each time point by dividing the signal measured in drug conditions with the initial luminescence of the untreated control. The minimum duration for killing (MDK) for each strain was determined by using the resulting killing data as input for in our “get MDK” script. MDK thresholds were chosen to maximise the number of strains that could be included in downstream analyses. For Mfx-treated strains, a threshold of 90% was used due to a strain exhibiting extensively high tolerance compared to others, whereas a threshold of 98% was used for RIF-treated strains.

### Statistical analysis

Statistical analysis and data visualization were performed using GraphPad Prism version 8.2.1. Tolerance and MIC data were imported into the Prism program and visualized. Data were tested for normality and t-tests were used to compare data between groups; otherwise, non-parametric equivalents were used. Specific statistical tests are specified in figure captions.

## Results and Discussion

### A cohort of TB patients with diverse clinical outcomes

We looked at retrospective laboratory clinical data from Georgian active pulmonary TB patients. We selected patients with multiple serial Mtb isolates with phenotypic drug susceptibility data that fit into one of four groups: DS-TB control (Group A, n = 3), DS-TB recalcitrant (Group B n= 6), MDR-TB control (Group C, n= 4) and MDR-TB resistance amplification (Group D, n = 5). All patients were diagnosed, monitored, and treated according to the Georgian National TB Program which includes molecular and culture-based drug susceptibility testing (30). Group A patients were initially diagnosed with DS-TB and became culture-negative within 12 months of initiating treatment. Group B patients were also diagnosed as DS-TB but remained sputum culture-positive for up to six years despite receiving treatment, without drug resistance evolving. Group C patients were diagnosed with MDR-TB and remained classified as such throughout treatment. Group D patients were diagnosed with MDR-TB and gained additional resistance during the course of treatment. This cohort, comprised of 18 patients and 28 strains, presented an interesting dataset in which to interrogate the clinical relevance of antibiotic tolerance in TB (**Fig 1, Table 1**).

**Table 1.** Strain information.

| Strain | Patient | Isolate #<br>/Total | Timespan<br>of<br>isolates<br>(Months) | DST | Lineage | Category |
| --- | --- | --- | --- | --- | --- | --- |
| N50209 | P01 | 1/8 | 56.13333 | Susceptible | L2 | Group B |
| N50215 | P01 | 8/8 |  | Susceptible | L2 | Group B |
| N50225 | P02 | 1/9 | 75.13333 | Susceptible | L4 | Group B |
| N50232 | P02 | 8/9 |  | Susceptible | L4 | Group B |
| N50247 | P03 | 1/8 | 52.46667 | Susceptible | L4 | Group B |
| N50255 | P03 | 8/8 |  | Susceptible | L4 | Group B |
| N50323 | P04 | 1/5 | 79.23333 | Susceptible | L4 | Group B |
| N50327 | P04 | 5/5 |  | Susceptible | L4 | Group B |
| N50339 | P05 | 1/9 | 22.13333 | Susceptible | L4 | Group B |
| N50246 | P05 | 9/9 |  | Susceptible | L4 | Group B |
| N50183 | P06 | 1/8 | 53.53333 | Susceptible | L4 | Group B |
| N50189 | P06 | 7/8 |  | Susceptible | L4 | Group B |
| N50445 | P07 | 1/4 | 12.83333 | Susceptible | L4 | Group A |
| N50447 | P08 | 1/4 | 2.13333 | Susceptible | L4 | Group A |
| N50450 | P09 | 1/4 | 3.9 | Susceptible | L4 | Group A |
| N50180 | P10 | 1/4 | 7.5 | MDR | L2 | Group C |
| N50182 | P10 | 4/4 |  | MDR | L2 | Group C |
| N50262 | P11 | 1/5 | 14 | MDR | L2 | Group C |
| N50265 | P11 | 5/5 |  | MDR | L2 | Group C |
| N50335 | P12 | 1/4 | 8.73333 | MDR | L4 | Group C |
| N50338 | P12 | 4/4 |  | MDR | L4 | Group C |
| N50238 | P13 | 1/6 | 81.56667 | MDR | L2 | Group C |
| N50241 | P13 | 4/6 |  | MDR | L2 | Group C |
| N50205 | P14 | 1/4 | 31.03333 | MDR | L2 | Group D |
| N50267 | P15 | 2/3 | 1.86667 | MDR | L2 | Group D |
| N50269 | P16 | 1/5 | 17.1 | MDR | L2 | Group D |
| N50233 | P17 | 1/4 | 5.06667 | MDR | L2 | Group D |
| N50278 | P18 | 1/5 | 12.46667 | MDR | L2 | Group D |

**Figure 1.**
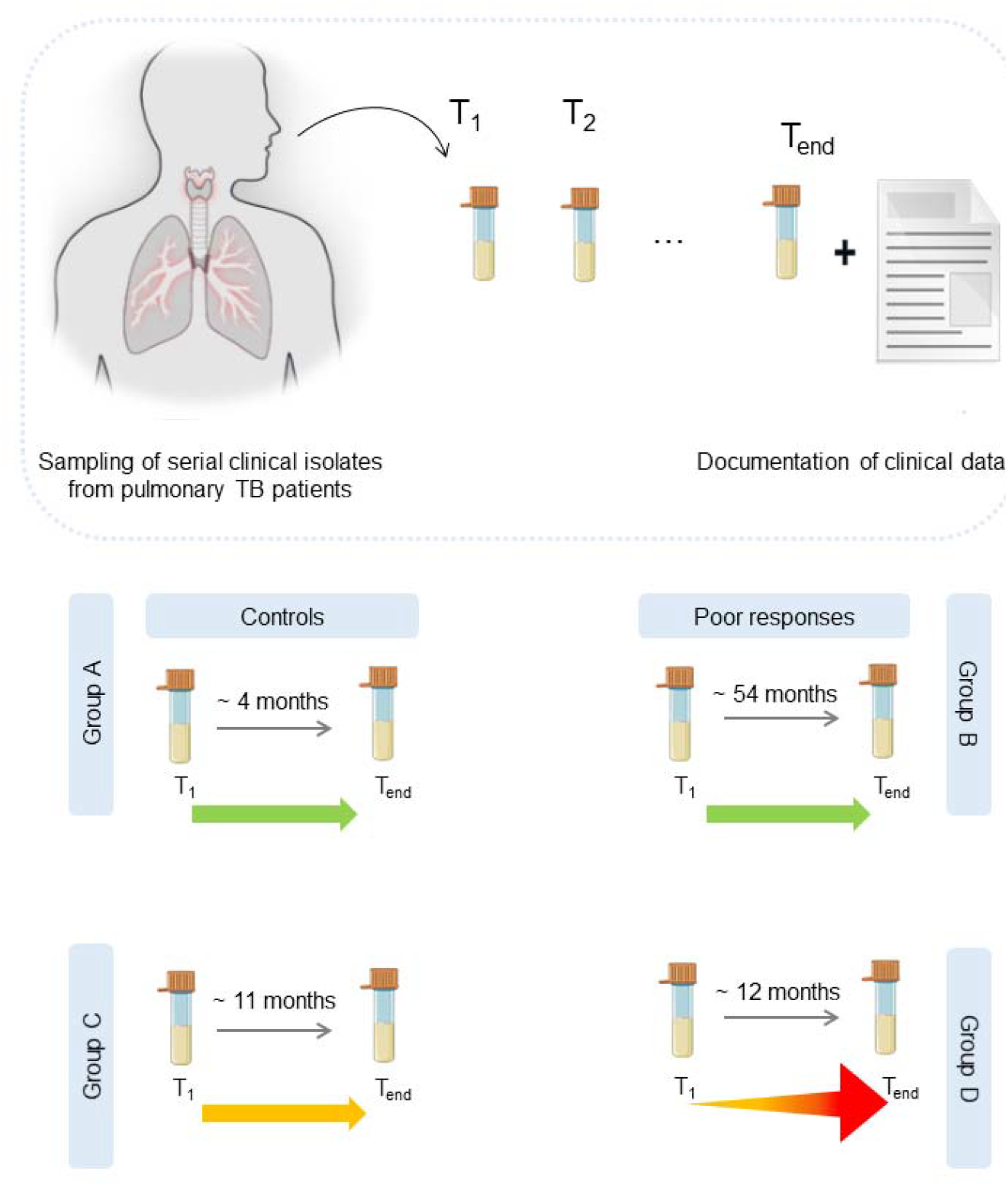
clinical Scheme of isolate origin and strain classification. Serial Mtb isolates were collected from TB patients throughout their TB episodes. Initial (T_1_) and final (T_end_) isolates from patients were selected and categorized into groups according to drug susceptibility and patient responses. Time span between isolates in diagram represent median time span of isolates within the group. Group A strains (n =3 strains and patients) were phenotypically drug susceptible and were cured within 13 months. Group B strains (n= 12) were drug susceptible and remained so, but patients (n = 6) had delayed culture conversion. Group C strains (n =8 strains, n = 4 patients) were MDR and remained so until final sampling. Group D strains (n = 5) were initially MDR and patients (n = 5) experienced drug resistance amplification.

To confirm that patients were infected with the same genotype throughout the course of their episode and to genotypically monitor drug resistance markers, we performed whole genome analysis on all isolates. From our phylogenetic analysis, we saw that isolates from the same patient separated in time clustered together with a SNP distance less than a maximum of 16 SNPs, supporting the notion that they were effectively the same strain (**Fig S1, Table S1**). MDR isolates were mainly members of *Mycobacterium tuberculosis* complex (MTBC) lineage (L) 2, whereas drug-susceptible isolates were mainly members of L4 which corresponds with general Georgian TB molecular epidemiology (25, 31, 32); as such, our sample set was endemically relevant.

### Tolerance does not explain poor treatment responses

While there has been evidence that tolerance exists and varies in Mtb (33, 34), its clinical relevance has not been thoroughly established. To investigate whether Group B cases were more tolerant to RIF and thus remained culture positive without gaining resistance, we measured RIF tolerance in Group A cases compared to Group B cases. We found no significant differences in RIF MDK_98_ between strains from group A and Group B patients (**Fig 2A**). In addition, RIF MICs were relatively stable (**Fig S2**), thus RIF susceptibility could not explain recalcitrant TB in our strain set.

**Figure 2.**
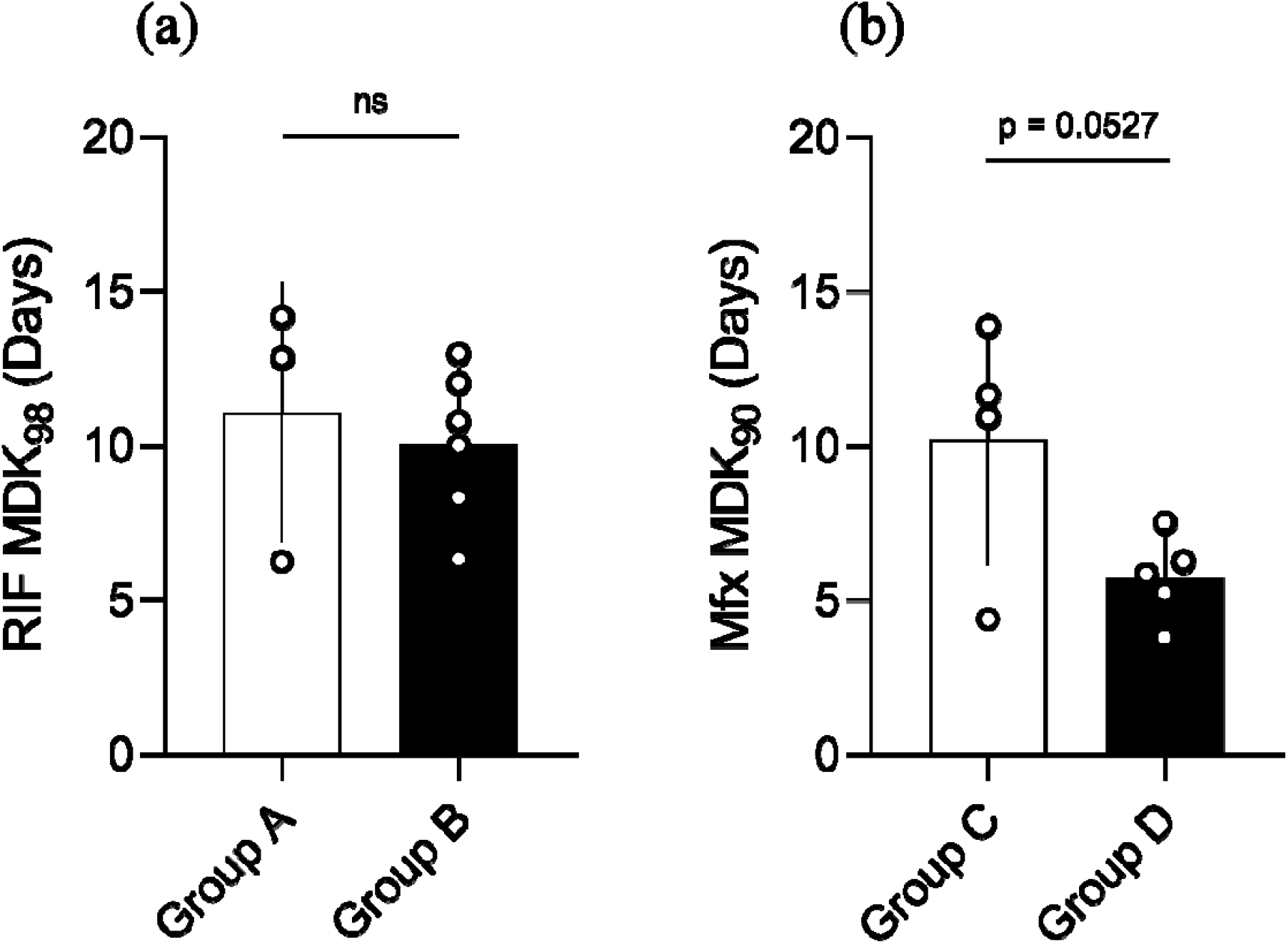
Differences in RIF tolerance do not account for recalcitrant TB, but low Mfx tolerance may play a role in resistance amplification. **A.** Comparison of RIF tolerance in strains isolated from control vs recalcitrant susceptible patients. Bars denote mean, error bars indicate standard deviation, ns = not significant by unpaired t-test. **B**. Comparison of tolerance to Mfx in strains isolated from control vs resistance amplification patients. Bars denote mean, error bars indicate standard deviation, p =0.0527 by unpaired t-test.

To explore whether Mfx tolerance is involved in resistance amplification in MDR-TB, we compared Mfx tolerance between Group C and Group D strains. Contrary to what has previously been reported in other bacteria, we saw lower Mfx MDK_90_ values in Group D strains compared to Group C. While this difference did not reach formal statistical significance (**Fig 2B**, p = 0.0527), the strains that eventually gained fluoroquinolone resistance to become pre-XDR by definition (6), had notably lower Mfx tolerance compared to those that remained MDR. Of note, the median time interval between initial and final isolate between group C and D patients was similar (~11 and 12 months, **Fig 1**), showing that within the same treatment window, some strains experienced resistance amplification while others did not.

In other bacteria, tolerance has been shown to lead to resistance evolution (27–29). Specifically, if two strains with equal drug susceptibility undergo a protocol of high-dose intermittent exposure to antibiotics, the strain that started more tolerant would more readily evolve drug resistance. In our data, lower Mfx tolerance was associated with resistance amplification, which goes against this convention and requires further exploration.

### Within-host evolution of tolerance has different trajectories

TB treatment requires a minimum of 6 months of antibiotic chemotherapy, which allows for within-host adaption of Mtb. Serial isolates provide an opportunity to observe such adaption across time. Having analysed the initial and final isolates from Group B and Group C patients, we were able to evaluate the dynamics of tolerance across time within-host.

RIF tolerance was heterogeneous across isolates from Group B patients, where tolerance increased (P03, P05, P06), remained stable (P01 and P04) or decreased (P04) across time (**Fig 3A**). The direction of the change in RIF tolerance did not appear to be related to the time interval between isolates (**Table 1**). The increase in RIF tolerance was notably higher by ~1.6-fold and ~1.3-fold in P03 and P06 final isolates, respectively, compared to their initial isolates (**Fig 3A**). This could indicate that higher RIF tolerance could be an effect of delayed culture conversion rather than a cause as we had initially expected, or as reported in other bacteria (29). This could be due to poor treatment adherence or insufficient treatment as in patients P03 and P06 who are both retreatment cases, thus representing intermittent drug exposure for the bacteria inside the host. *In vitro*, intermittent exposure to antibiotics evolves drug tolerance (27, 29), as such, intermittent treatment is a plausible contributing factor to the evolution of drug tolerance in TB patients.

**Figure 3.**
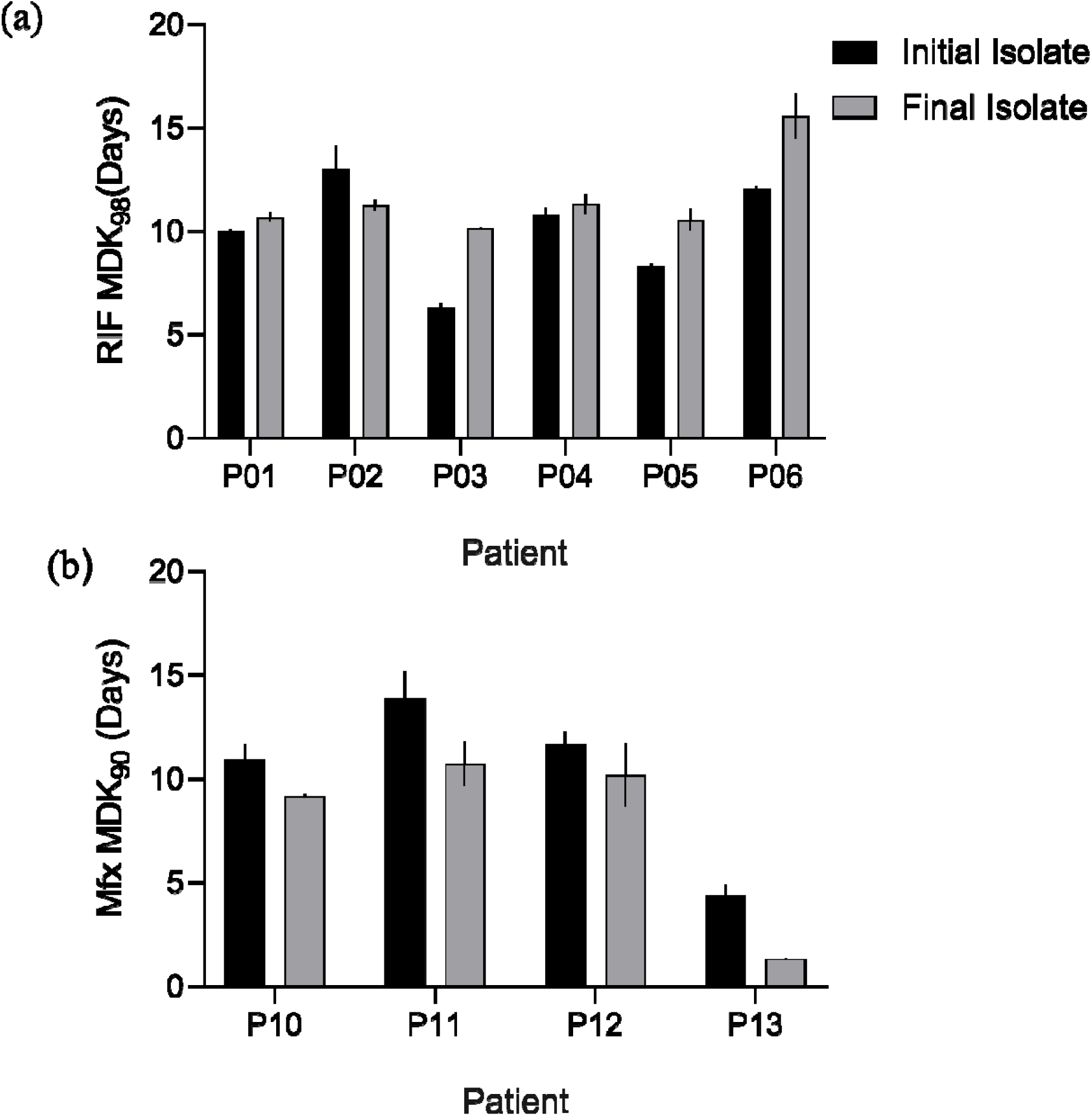
Within-host of dynamic of tolerance in TB patients. **A.** Comparison of change in tolerance between initial and final isolates from Group B patients. **B**. Comparison of changes in tolerance in Group C patients. Bars denote a mean of n = 2 independent experiments; error bars denote standard error.

In Group C, final isolates consistently exhibited lower Mfx MDK_90_ values and thus tolerance as compared to the initial isolate, both within host (**Fig 3B**), and among hosts (**Fig S6**). This finding indicates a tendency for Mfx tolerance to decrease over time in Mtb in patients undergoing MDR-TB treatment. This subtle evidence of a reversal of drug tolerance over time is unprecedented, but the decrease was consistent among all patients, stoking a need for further investigation.

## Concluding Remarks

Here we explored antibiotic tolerance in Mtb isolates from TB patients with defined treatment responses with respect to recalcitrance and resistance amplification. Our work has several limitations. First, our patient sample sizes are small. Despite this, we have been able to observe different manifestations of the dynamics of within-host drug tolerance. Second, due to the age of some of our Mtb isolates, some patient data was irretrievable. More complete clinical data could shed further light on where these dynamic changes in tolerance clinically manifest. Finally, while we set out to understand bacterial determinants that contribute to poor outcomes, we neglected to consider patient features within the drug-pathogen interaction. Patients can differ in their metabolism of TB drugs (35, 36), which could result in bacteria experiencing sub-lethal doses, thereby influencing culture conversion and acquired drug resistance (37). While we did not have access to patient pharmacokinetics and pharmacodynamics, which also affect treatment efficacy (38–40), in our historical data, future prospective studies endeavouring to understand within-host drug-shaped Mtb evolution, should consider this.

In conclusion, we provide evidence that antibiotic tolerance changes within-host in a patient and drug-specific manner, with an unclear contribution to patient treatment outcomes. However, it is likely that drug susceptibility by way of drug tolerance, alongside host-derived features work in concert in contributing to treatment responses, and as such should be further evaluated in larger sample sets.

## Acknowledgements

Calculations were performed at sciCORE (http://scicore.unibas.ch/) scientific computing core facility at the University of Basel. Sequencing was carried out at the Genomics Facility Basel of the University of Basel and the Department of Biosystems Science and Engineering at ETHZ in Basel, Switzerland. Credit to servier for screwcap tube icon, liquid colouring was altered.(tube-screwcap-closed-orange icon by Servier https://smart.servier.com/ is licensed under CC-BY 3.0 Unported https://creativecommons.org/licenses/by/3.0/).

## Transparency declarations

All authors have no conflict of interests to declare.

## Author Contributions

Study conception: V.F.A.M., L.J., N.T., G.A.G., S.Bor., and S.G; experimental design: V.F.A.M., L.J., S.Bor., and S.G.; data acquisition and analysis: V.F.A.M., K.M., N.M., T.A., A.D., S.Bou., and L.J.; interpretation of data: V.F.A.M., S.Bor., and S.G.; drafted the main manuscript text: V.F.A.M., S.Bor., and S.G.

## References

1. WHO. 2024. Global tuberculosis report 2024. World Health Organization.

2. Chan K-PF, Ting-Fung M, Sridhar S, Lui MM-S, Ho JC-M, Lam DC-L, Ip MS-M, Ho P-L. 2024. Changes in the incidence, clinical features and outcomes of tuberculosis during COVID-19 pandemic. Journal of Infection and Public Health 17:102511.

3. Jnawali HN, Ryoo S. 2013. First-and second-line drugs and drug resistance. Tuberculosis-current issues in diagnosis and management 20:163–80.

4. Ramachandran G, Swaminathan S. 2015. Safety and tolerability profile of second-line anti-tuberculosis medications. Drug safety 38:253–269.

5. Suárez PG, Floyd K, Portocarrero J, Alarcón E, Rapiti E, Ramos G, Bonilla C, Sabogal I, Aranda I, Dye C. 2002. Feasibility and cost-effectiveness of standardised second-line drug treatment for chronic tuberculosis patients: a national cohort study in Peru. The Lancet 359:1980–1989.

6. Organization WH. 2022. WHO consolidated guidelines on tuberculosis. Module 4: treatment-drug-resistant tuberculosis treatment, 2022 update. World Health Organization.

7. Conradie F, Bagdasaryan TR, Borisov S, Howell P, Mikiashvili L, Ngubane N, Samoilova A, Skornykova S, Tudor E, Variava E. 2022. Bedaquiline–pretomanid–linezolid regimens for drug-resistant tuberculosis. New England Journal of Medicine 387:810–823.

8. Ndjeka N, Schnippel K, Master I, Meintjes G, Maartens G, Romero R, Padanilam X, Enwerem M, Chotoo S, Singh N. 2018. High treatment success rate for multidrug-resistant and extensively drug-resistant tuberculosis using a bedaquiline-containing treatment regimen. European Respiratory Journal 52.

9. Goig GA, Loiseau C, Maghradze N, Mchedlishvili K, Avaliani T, Brites D, Borell S, Aspindzelashvili R, Avaliani Z, Kipiani M. 2024. Transmission is a key driver of extensively drug-resistant tuberculosis. medRxiv:2024.06.28.24309543.

10. Atif M, Anwar Z, Fatima RK, Malik I, Asghar S, Scahill S. 2018. Analysis of tuberculosis treatment outcomes among pulmonary tuberculosis patients in Bahawalpur, Pakistan. BMC Research Notes 11:370.

11. Fekadu G, Turi E, Kasu T, Bekele F, Chelkeba L, Tolossa T, Labata BG, Dugassa D, Fetensa G, Diriba DC. 2020. Impact of HIV status and predictors of successful treatment outcomes among tuberculosis patients: A six-year retrospective cohort study. Annals of Medicine and Surgery 60:531–541.

12. Jackson C, Stagg H, Doshi A, Pan D, Sinha A, Batra R, Batra S, Abubakar I, Lipman M. 2017. Tuberculosis treatment outcomes among disadvantaged patients in India. Public Health Action 7:134–140.

13. Khaliaukin A, Kumar A, Skrahina A, Hurevich H, Rusovich V, Gadoev J, Falzon D, Khogali M, De Colombani P. 2014. Poor treatment outcomes among multidrug-resistant tuberculosis patients in Gomel Region, Republic of Belarus. Public Health Action 4:S24–S28.

14. Muñoz-Sellart M, Cuevas L, Tumato M, Merid Y, Yassin M. 2010. Factors associated with poor tuberculosis treatment outcome in the Southern Region of Ethiopia. The International Journal of Tuberculosis and Lung Disease 14:973–979.

15. Teferi MY, El-Khatib Z, Boltena MT, Andualem AT, Asamoah BO, Biru M, Adane HT. 2021. Tuberculosis Treatment Outcome and Predictors in Africa: A Systematic Review and Meta-Analysis. International Journal of Environmental Research and Public Health 18:10678.

16. Vasankari T, Holmström P, Ollgren J, Liippo K, Kokki M, Ruutu P. 2007. Risk factors for poor tuberculosis treatment outcome in Finland: a cohort study. BMC Public Health 7:291.

17. Alemu A, Bitew ZW, Worku T. 2020. Poor treatment outcome and its predictors among drug-resistant tuberculosis patients in Ethiopia: A systematic review and meta-analysis. International Journal of Infectious Diseases 98:420–439.

18. Nair D, Velayutham B, Kannan T, Tripathy J, Harries A, Natrajan M, Swaminathan S. 2017. Predictors of unfavourable treatment outcome in patients with multidrug-resistant tuberculosis in India. Public Health Action 7:32–38.

19. Wen Y, Zhang Z, Li X, Xia D, Ma J, Dong Y, Zhang X. 2018. Treatment outcomes and factors affecting unsuccessful outcome among new pulmonary smear positive and negative tuberculosis patients in Anqing, China: a retrospective study. BMC Infectious Diseases 18:104.

20. García-Basteiro AL, Respeito D, Augusto OJ, López-Varela E, Sacoor C, Sequera VG, Casellas A, Bassat Q, Manhiça I, Macete E, Cobelens F, Alonso PL. 2016. Poor tuberculosis treatment outcomes in Southern Mozambique (2011–2012). BMC Infectious Diseases 16:214.

21. Ifebunandu NA, Ukwaja KN, Obi SN. 2012. Treatment outcome of HIV-associated tuberculosis in a resource-poor setting. Tropical Doctor 42:74–76.

22. Tang S, Tan S, Yao L, Li F, Li L, Guo X, Liu Y, Hao X, Li Y, Ding X. 2013. Risk factors for poor treatment outcomes in patients with MDR-TB and XDR-TB in China: retrospective multi-center investigation. PLoS one 8:e82943.

23. Javaid A, Ullah I, Masud H, Basit A, Ahmad W, Butt ZA, Qasim M. 2018. Predictors of poor treatment outcomes in multidrug-resistant tuberculosis patients: a retrospective cohort study. Clinical Microbiology and Infection 24:612–617.

24. Johnston JC, Shahidi NC, Sadatsafavi M, Fitzgerald JM. 2009. Treatment outcomes of multidrug-resistant tuberculosis: a systematic review and meta-analysis. PloS one 4:e6914.

25. Goig GA, Loiseau C, Maghradze N, Mchedlishvili K, Avaliani T, Tsutsunava A, Brites D, Kalkan S, Borrell S, Aspindzelashvili R. 2025. Clinical and bacterial determinants of unfavorable tuberculosis treatment outcomes: an observational study in Georgia. medRxiv:2025.01.20.25320828.

26. Brauner A, Fridman O, Gefen O, Balaban NQ. 2016. Distinguishing between resistance, tolerance and persistence to antibiotic treatment. Nature Reviews Microbiology 14:320–330.

27. Santi I, Manfredi P, Maffei E, Egli A, Jenal U. 2021. Evolution of antibiotic tolerance shapes resistance development in chronic Pseudomonas aeruginosa infections. MBio 12:10.1128/mbio.03482-20.

28. Levin-Reisman I, Ronin I, Gefen O, Braniss I, Shoresh N, Balaban NQ. 2017. Antibiotic tolerance facilitates the evolution of resistance. Science 355:826–830.

29. Liu J, Gefen O, Ronin I, Bar-Meir M, Balaban NQ. 2020. Effect of tolerance on the evolution of antibiotic resistance under drug combinations. Science 367:200–204.

30. March VF, Zwyer M, Loiseau C, Brites D, Goig GA, Bouaouina S, Dötsch A, Reinhardt M, Kalkan S, Gagneux S. 2025. Variability in intrinsic drug tolerance in Mycobacterium tuberculosis corresponds with phylogenetic lineage. bioRxiv:2025.07.04.663131.

31. Gygli SM, Loiseau C, Jugheli L, Adamia N, Trauner A, Reinhard M, Ross A, Borrell S, Aspindzelashvili R, Maghradze N. 2021. Prisons as ecological drivers of fitness-compensated multidrug-resistant Mycobacterium tuberculosis. Nature medicine 27:1171–1177.

32. Loiseau C, Windels EM, Gygli SM, Jugheli L, Maghradze N, Brites D, Ross A, Goig G, Reinhard M, Borrell S. 2023. The relative transmission fitness of multidrug-resistant Mycobacterium tuberculosis in a drug resistance hotspot. Nature communications 14:1988.

33. Vijay S, Nhung HN, Bao NLH, Thu DDA, Trieu LPT, Phu NH, Thwaites GE, Javid B, Thuong NT. 2021. Most-probable-number-based minimum duration of killing assay for determining the spectrum of rifampicin susceptibility in clinical Mycobacterium tuberculosis isolates. Antimicrobial Agents and Chemotherapy 65:10.1128/aac.01439-20.

34. Srinivasan V, Bao NLH, Vinh DN, Le Quang N, Nhung HN, Ha VTN, Thai PVK, Ha DTM, Lan NH, Caws M. 2024. Rifampicin tolerance and growth fitness among isoniazid-resistant clinical Mycobacterium tuberculosis isolates: an in-vitro longitudinal study. eLife 13.

35. Chideya S, Winston CA, Peloquin CA, Bradford WZ, Hopewell PC, Wells CD, Reingold AL, Kenyon TA, Moeti TL, Tappero JW. 2009. Isoniazid, rifampin, ethambutol, and pyrazinamide pharmacokinetics and treatment outcomes among a predominantly HIV-infected cohort of adults with tuberculosis from Botswana. Clinical infectious diseases 48:1685–1694.

36. Sileshi T, Tadesse E, Makonnen E, Aklillu E. 2021. The Impact of First-Line Anti-Tubercular Drugs’ Pharmacokinetics on Treatment Outcome: A Systematic Review. Clinical Pharmacology: Advances and Applications 13:1–12.

37. Pasipanodya JG, Srivastava S, Gumbo T. 2012. Meta-analysis of clinical studies supports the pharmacokinetic variability hypothesis for acquired drug resistance and failure of antituberculosis therapy. Clinical Infectious Diseases 55:169–177.

38. Liu Q, Wei J, Li Y, Wang M, Su J, Lu Y, López MG, Qian X, Zhu Z, Wang H. 2020. Mycobacterium tuberculosis clinical isolates carry mutational signatures of host immune environments. Science advances 6:eaba4901.

39. Alffenaar J-WC, Gumbo T, Dooley KE, Peloquin CA, Mcilleron H, Zagorski A, Cirillo DM, Heysell SK, Silva DR, Migliori GB. 2020. Integrating pharmacokinetics and pharmacodynamics in operational research to end tuberculosis. Clinical Infectious Diseases 70:1774–1780.

40. Munro SA, Lewin SA, Smith HJ, Engel ME, Fretheim A, Volmink J. 2007. Patient adherence to tuberculosis treatment: a systematic review of qualitative research. PLoS medicine 4:e238.

