## Supplemental1 for "Within-host evolution of drug tolerance in *Mycobacterium tuberculosis*"

**Supplementary information**

**Methods:**

**Strain stock preparation**

Serial Mtb isolates were sampled from these patients as part of the Georgian National TB Control Program, and according to their regulations as previously described (1). Briefly, patient sputa were collected and decontaminated using the NaOH approach. Decontaminated samples were processed for smear microscopy, and drug susceptibility testing using GeneXpert, line probe assays and the MGIT BACTEC 960^TM^ system. Isolates were frozen from turbid MGIT cultures by transferring 500-800 µl from the bottom of the MGIT tubes into cryovials containing 50% glycerol diluted in phosphate buffer, for a final concentration of 5% glycerol. Stocks were frozen at - 80 °C.

Based on clinical drug susceptibility and genomic data, strains from the biobank at NCTLD were selected and shipped to our BSL3 facilities in Switzerland at room temperature and then stored in our - 80 °C freezer. Working stocks were prepared from the imported strains by inoculating at least 20 µl of original stocks in 7H9 ADC. Strains were incubated at 37 °C with shaking at 140 rpm until they reached log to early stationary phase (OD_600nm_ 0.8 – 1). When ready, strains were centrifuged at 800 *x g* and the supernatant was discarded. Strains were resuspended in volumes of 2 ml in 7H9 ADC + 5% glycerol (PanReac AppliedChem) and frozen in 250 µl aliquots. Strains were routinely cultured in 7H9 ADC at 37 °C shaking at 140 rpm. Calibrated stocks of working strains were prepared as previously described (**Chapter 5**), briefly, strains were grown to mid-log phase (OD_600nm_ 0.4 – 0.8), and clumps were pelleted by centrifugation at 260 x g. The resulting supernatant was calibrated in 7H9 ADC to an OD_600nm_ of 0.1 and frozen at - 80 °C in 1 ml aliquots.

**DNA Extraction and whole genome sequencing (WGS)**

DNA was extracted in Georgia from selected patient strains. Strains were grown in 7H9 broth supplemented with 10% ADC (5% bovine albumin-fraction V + 2% dextrose + 0.003% catalase, Sigma-Aldrich), 0.5% glycerol (PanReac AppliedChem) and 0.1% Tween-80 (Sigma-Aldrich), hence 7H9 ADC until cultures were highly turbid, and DNA was extracted using the CTAB method(2). DNAs were shipped to Switzerland and sent to the Department of Department of Biosystems Science and Engineering ETH Zurich in Basel for library preparation and whole genome sequencing using Illumina short-read technology.

**WGS and phylogenetic analyses:**

The sequences used for analysis in this study include previously sequenced and newly sequenced strains. Accession numbers are detailed in **supplementary** **data**.

FastQ files were analysed using our in-house pipeline previously described (3). In summary, sequencing reads were taxonomically classified and identified as belonging to the Mycobacterium tuberculosis complex (MTBC) using kraken v.1.1.1 (4). Illumina adapters were removed and reads were trimmed using Trimmomatic (5) v0.39 using a 5 bp sliding window (cutting off when the median quality dropped below 20). Resulting reads shorter than 20 bp were discarded. For paired-end data, SeqPrep v1.3.1 (https://github.com/jstjohn/SeqPrep) was employed to merge overlapping reads, provided the overlap was at least 15 bp. Prior to mapping, reads classified by Kraken as non-MTBC were removed. The resulting high-quality MTBC reads were aligned to the inferred ancestor of the MTBC (6) using BWA v0.7.17 (10.5281/zenodo.3497110) (7). Duplicate reads were flagged using the MarkDuplicates module of Picard v2.26.2 (<http://broadinstitute.github.io/picard/>). Variant calling was performed using the mutect2 module of GATK v4.2.3 (8) and then variants were filtered using FilterMutectCalls in microbial mode. Variants were annotated and their effects on genes was predicted using SnpEff v5.0 (9). We excluded variants in repetitive regions such as PE, PPE, and PGRS genes or phages, as decribed in Stucki et al 2016 (10).

Drug resistance-conferring variants were annotated using the second version of the WHO catalogue and only mutations having a frequency >= 90% were kept for the analysis. Strain drug susceptibility was based on the presence of drug resistance conferring mutations in targets of key drugs as per the WHO 2022 definitions. Lineages were identified based on single nucleotide polymorphisms as described in Coll et al 2014 (11).

A multiple sequence alignment using quality filtered fasta files was generated as previously described(12) from patient strain genomes, retaining drug resistance variants. A maximum likelihood phylogeny was generated using IQ-TREE version 2.1.3(13) with *Mycobacterium canettii* as an outgroup. Analysis was run with 1000 pseudo-replicates in ultrafast bootstrap mode, which was visualized and annotated using interactive Tree of Life (iTOL) version 7(14).

To calculate single nucleotide polymorphism (SNP) distances between isolates, we built an alignment of all non-redundant fixed SNPs observed across isolates of each patient and enumerated the number of fixed SNPs, defined by having a frequency >= 90%.

**Preparation of resazurin stocks**

Resazurin stocks were prepared by dissolving 3.125 mg of resazurin sodium salt (Sigma-Aldrich) in 25 ml of MilliQ water. After vortexing, the solution was sterilized in a sterile hood using a 0.22 μM syringe filter. Aliquots of the resulting resazurin solution were prepared at a volume of 1.1 ml, and frozen at -20 °C. For each use, aliquots were thawed overnight at 4 °C and equilibrated to room temperature before use.

**Measuring minimum inhibitory concentration (MIC)**

MICs of strains were determined using the REMA method as previously described(15-17). Briefly, 10 µl of thawed calibrated stocks (prepared as detailed above) were inoculated onto 96 well plates containing antibiotics in a 2-fold dilution series in 7H9 ADC across the plate with drug free wells for positive and negative controls. Each antibiotic concentration was inoculated in triplicate per strain and plates were incubated at 37 °C for 7 days for drug susceptible strains and 10 days for MDR strains. After incubation, 10 µl of resazurin was added to each well, and plates were incubated for a further 24 hours. Resorufin fluorescence was measured using the Tecan Infinite Pro 200, with excitation at 560 nm and emission measured at 590 nm. Relative growth was expressed as the ratio of the fluorescence of drug treated signal over the untreated control signal after subtracting background fluorescence The MIC was determined as the concentration at which growth was inhibited by at least 90%.

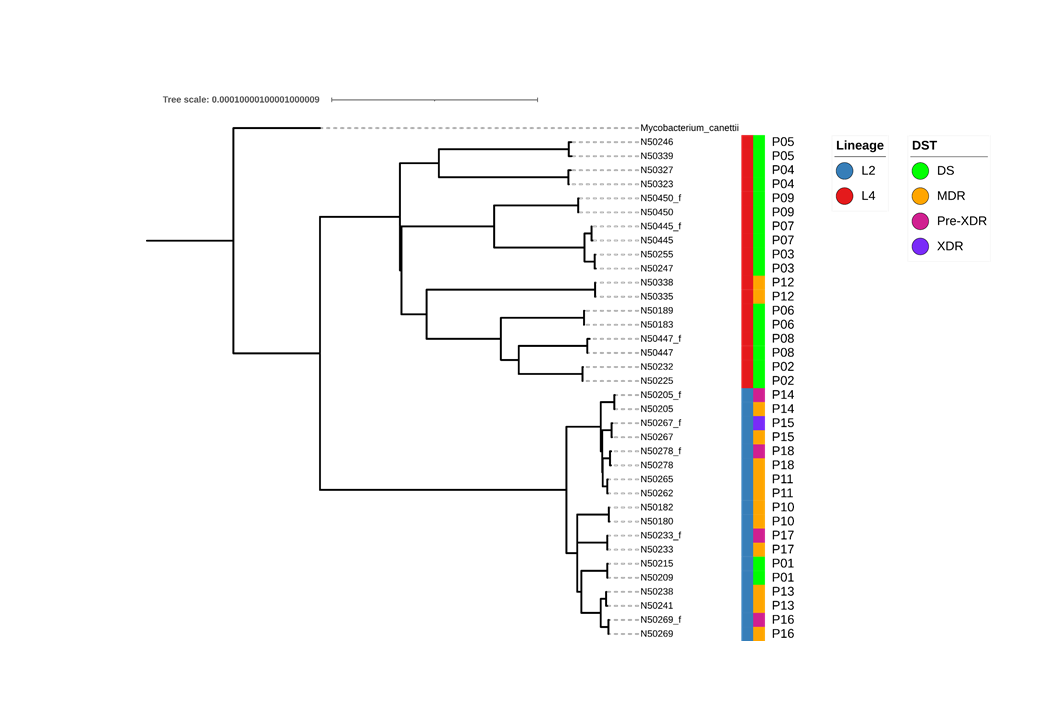
**Supplementary Information**

**Figure S1: Strain phylogeny.** Maximum likelihood tree of clinical strains sampled from historical TB patients from Georgia. Each imported strain has a unique identifier, while strains not imported include the suffix “f” to indicate that the genome comes from the final sample isolated from the patient. Tips labelled by strain identifier with lineage, DST, and patient number included. DST defined by pre-2022 WHO update.

**Table S1**: Single Nucleotide Polymorphism (SNP) distances between initial and final patient isolates.

| **Patient** | **SNP distance** |
| --- | --- |
| P01 | 7 |
| P02 | 9 |
| P03 | 10 |
| P04 | 9 |
| P05 | 9 |
| P06 | 4 |
| P07 | 3 |
| P08 | 7 |
| P09 | 2 |
| P10 | 3 |
| P11 | 4 |
| P12 | 2 |
| P13 | 16 |
| P14 | 3 |
| P15 | 4 |
| P16 | 4 |
| P17 | 6 |
| P18 | 5 |

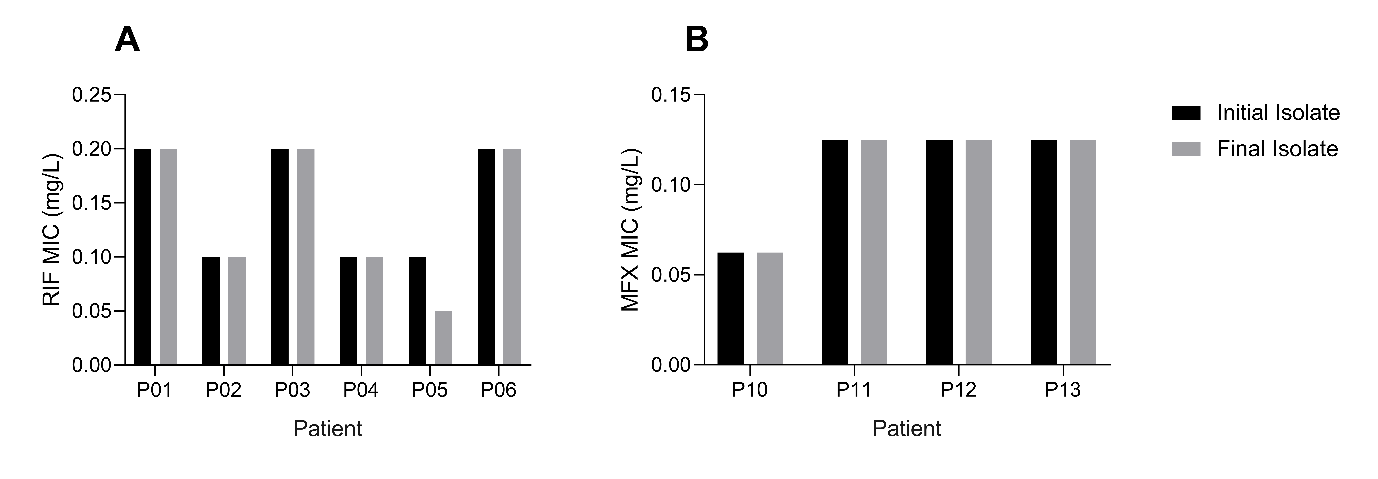

**Figure S2.** Relatively stable MICs across patient isolates. **A.** Rifampicin MICs for in initial and final isolates of sus-re patients. **B.** Moxifloxacin MICs for initial and final isolates of MDR-c patients.

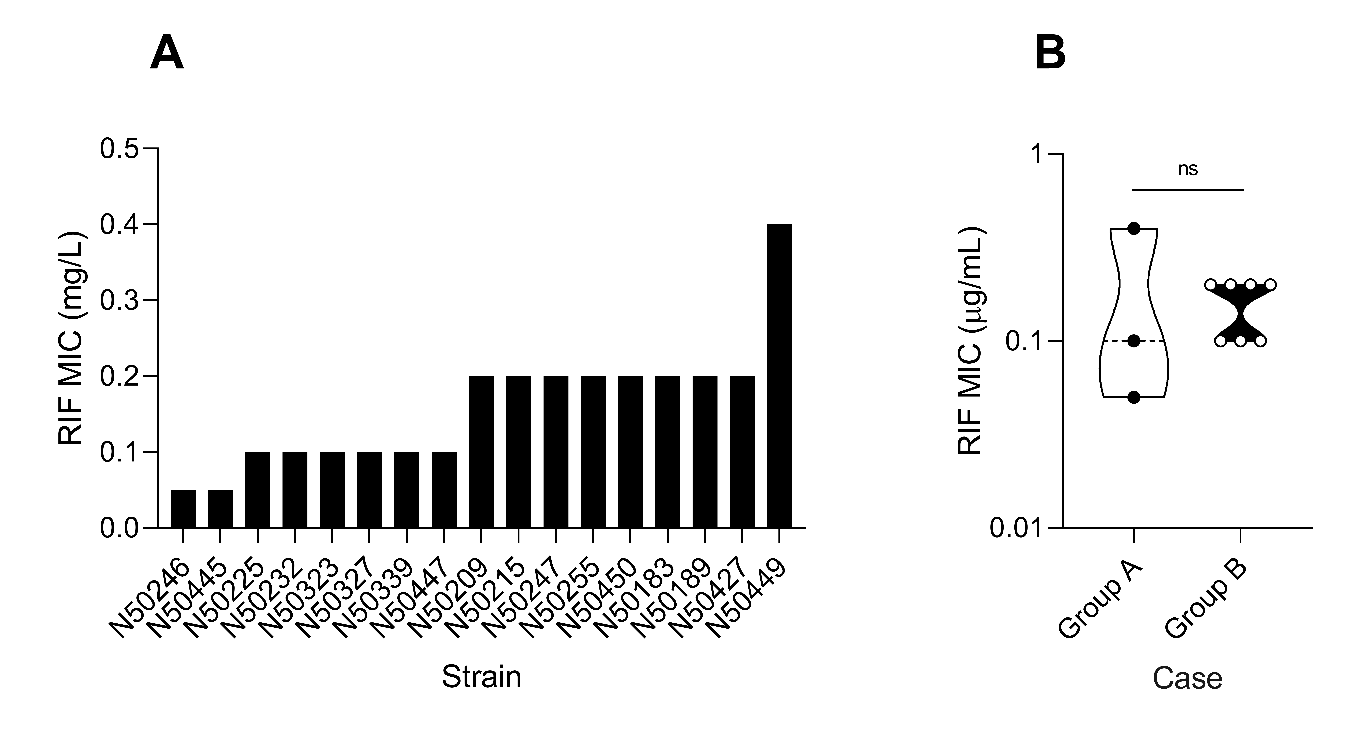

**Figure S3:** Rifampicin (RIF) susceptibility in clinical isolates **A.** Rifampicin MIC values for susceptible strains used in this study, values represent results of n = 2 independent experiments. **B.** Violin plots comparing strains originating from susceptible control (n = 3) patients and susceptible recalcitrant TB patients (n = 7), ns = not significant by unpaired t-test.

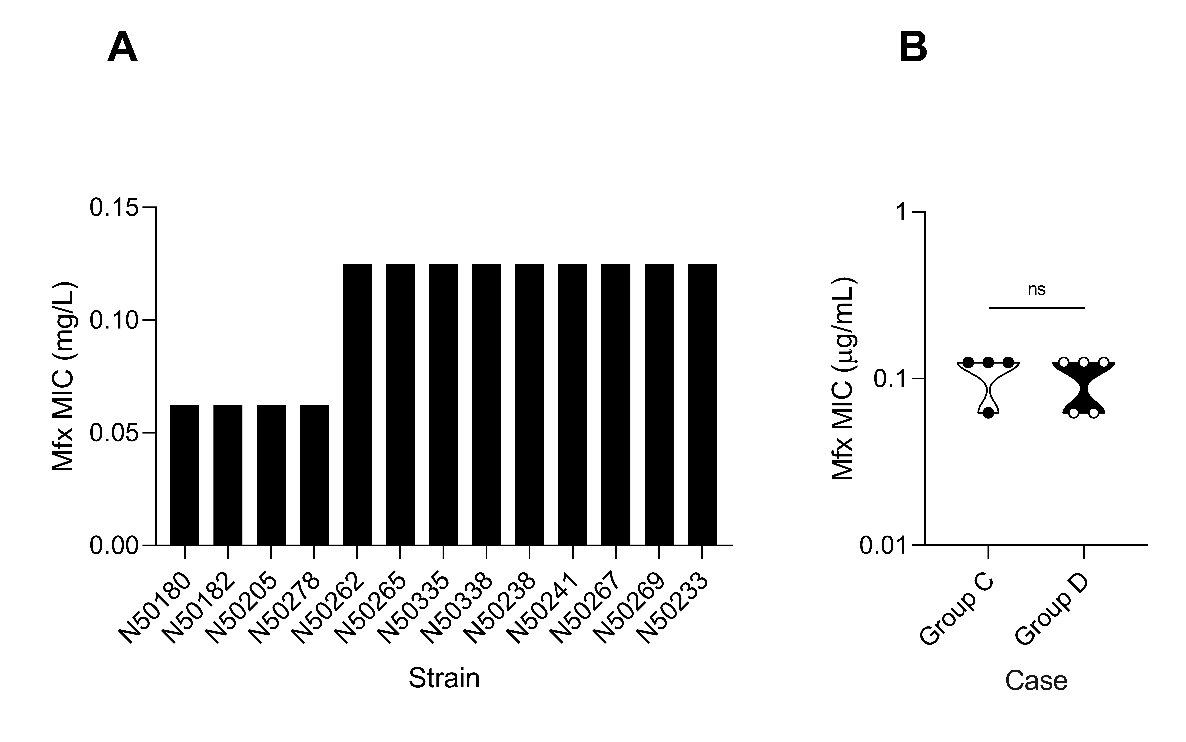

**Figure S4:** Moxifloxacin (Mfx) susceptibility in MDR clinical isolates. **A.** Moxifloxacin MIC of multidrug-resistant strains used in this study; data depicts MIC derived from two independent experiments. **B.** Comparison of MICs from MDR control (Group C, n = 4) patients and MDR resistance amplification (Group D, n = 5) patients, ns = not significant by unpaired t-test.

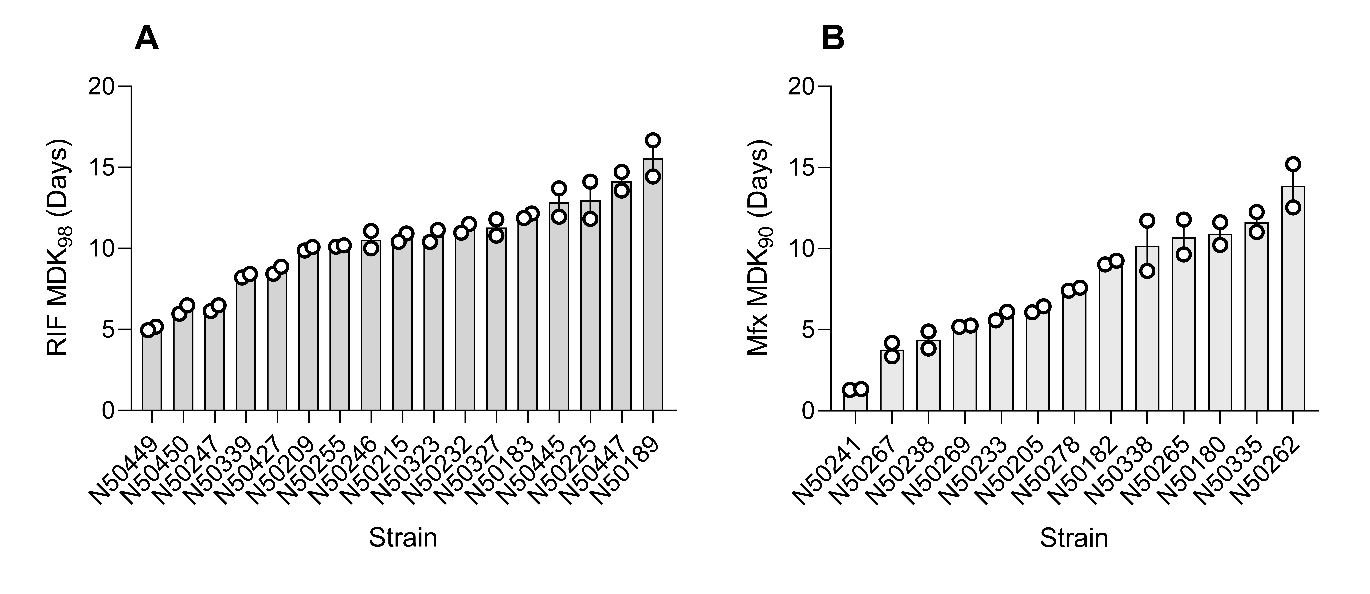

**Figure S5:** Variation in tolerance to RIF and Mfx**. A.** Bar graph depicting mean RIF tolerance in susceptible isolates data representative of n = 2 independent experiments, error bars depict standard error. **B.** Bar graphs depicting mean Mfx tolerance in MDR isolates, data representative of n = 2 independent experiments, error bars depict standard error.

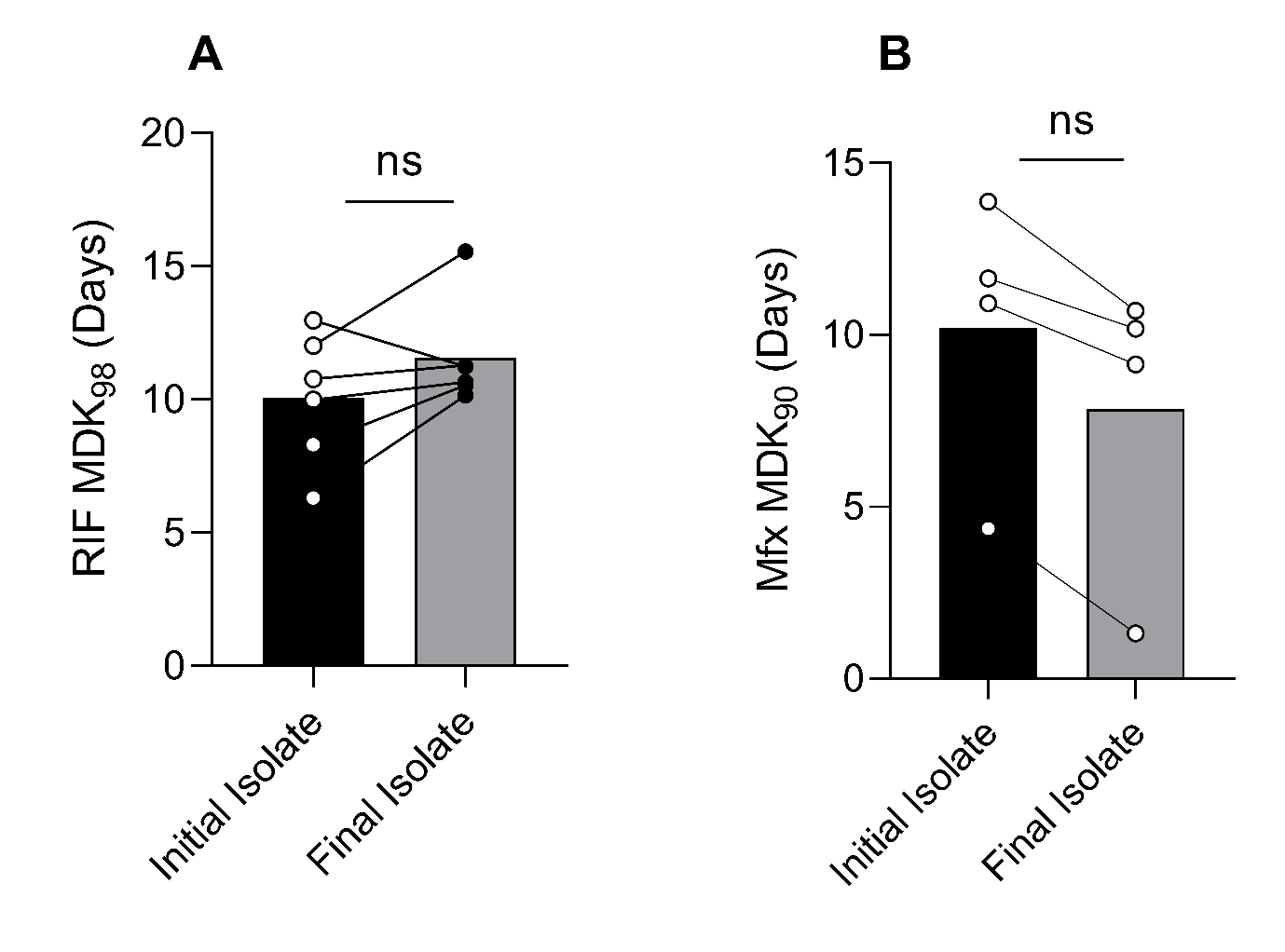

**Figure S6:** Global change in tolerance to RIF and Mfx in Group B and C patients respectively**. A.** Before and after plot of tolerance in strains from Group B patients. **B**. Before and after plot of tolerance in strains from Group C patients. Bar graphs depicting mean tolerance in group, dots represent mean of n = 2 independent experiments, ns = non-significant by Wilcoxon matched-pairs signed rank test.

| Category | Patient | Sex |  | Outcome | Age at Initial Diagnosis | Age at Outcome | ~Timespan of culture positivity (Months) | Case definition at diagnosis |
| --- | --- | --- | --- | --- | --- | --- | --- | --- |
| Group B | P01 | Male |  | Died | 42 | 47 | 56 | Treatment after default |
| Group B | P02 | Male |  | Default | 47 | 52 | 75 | New |
| Group B | P03 | Male |  | Default | 51 | 55 | 52 | Treatment after default |
| Group B | P04 | Male |  | Completed | 80 | 87 | 79 | Treatment after default |
| Group B | P05 | Male |  | Cured | 61 | 62 | 22 | New |
| Group B | P06 | Male |  | Died | 58 | 62 | 54 | Relapse |
| Group A | P07 | Male |  | Cured | 50 | 51 | 13 | New |
| Group A | P08 | Male |  | Cured | 49 | 49 | 2 | New |
| Group A | P09 | Male |  | Cured | 52 | 52 | 4 | New |

**Table S2:** Clinical information retrieved for susceptible TB patients

| **Category** | **Patient** | **Sex** | **Outcome** | **Age at Initial Diagnosis** | **Age at Outcome** | **~Timespan of culture positivity (Months)** | **Case definition at diagnosis** |
| --- | --- | --- | --- | --- | --- | --- | --- |
| Group C | P10 | NA | NA | NA | NA | 7.5 | NA |
| Group C | P11 | Male | Default | 37 | 38 | 14 | NA |
| Group C | P12 | NA | NA | NA | NA | 8.733333 | NA |
| Group C | P13 | Male | Default | NA | NA | 81.56667 | Treatment after Failure |
| Group D | P14 | NA | NA | NA | NA | 31.03333 | NA |
| Group D | P15 | Male | Cured | 49 | 49 | 1.866667 | New Case |
| Group D | P16 | Male | Died | 28 | 30 | 17.1 | Relapse |
| Group D | P16 | NA | NA | NA | NA | 5.066667 | NA |
| Group D | P18 | Male | Failure | 40 | 41 | 12.46667 | NA |

**Table S3.** Retrieved MDR patient clinical information

1.	March VF, Maghradze N, Mchedlishvili K, Avaliani T, Aspindzelashvili R, Avaliani Z, Kipiani M, Tukvadze N, Jugheli L, Bouaouina S. 2025. Drug-induced differential culturability in diverse strains of Mycobacterium tuberculosis. Scientific Reports 15:3588.

2.	Van Embden J, Cave MD, Crawford J, Dale J, Eisenach K, Gicquel B, Hermans P, Martin C, McAdam R, Shinnick T. 1993. Strain identification of Mycobacterium tuberculosis by DNA fingerprinting: recommendations for a standardized methodology. Journal of clinical microbiology 31:406-409.

3.	Goig GA, Loiseau C, Maghradze N, Mchedlishvili K, Avaliani T, Brites D, Borell S, Aspindzelashvili R, Avaliani Z, Kipiani M. 2024. Transmission is a key driver of extensively drug-resistant tuberculosis. medRxiv:2024.06. 28.24309543.

4.	Wood DE, Salzberg SL. 2014. Kraken: ultrafast metagenomic sequence classification using exact alignments. Genome Biology 15:R46.

5.	Bolger AM, Lohse M, Usadel B. 2014. Trimmomatic: a flexible trimmer for Illumina sequence data. Bioinformatics 30:2114-20.

6.	Comas I, Chakravartti J, Small PM, Galagan J, Niemann S, Kremer K, Ernst JD, Gagneux S. 2010. Human T cell epitopes of Mycobacterium tuberculosis are evolutionarily hyperconserved. Nature genetics 42:498-503.

7.	Li H, Durbin R. 2009. Fast and accurate short read alignment with Burrows–Wheeler transform. bioinformatics 25:1754-1760.

8.	McKenna A, Hanna M, Banks E, Sivachenko A, Cibulskis K, Kernytsky A, Garimella K, Altshuler D, Gabriel S, Daly M. 2010. The Genome Analysis Toolkit: a MapReduce framework for analyzing next-generation DNA sequencing data. Genome research 20:1297-1303.

9.	Cingolani P, Platts A, Wang LL, Coon M, Nguyen T, Wang L, Land SJ, Lu X, Ruden DM. 2012. A program for annotating and predicting the effects of single nucleotide polymorphisms, SnpEff: SNPs in the genome of Drosophila melanogaster strain w1118; iso-2; iso-3. fly 6:80-92.

10.	Stucki D, Brites D, Jeljeli L, Coscolla M, Liu Q, Trauner A, Fenner L, Rutaihwa L, Borrell S, Luo T. 2016. Mycobacterium tuberculosis lineage 4 comprises globally distributed and geographically restricted sublineages. Nature genetics 48:1535-1543.

11.	Coll F, McNerney R, Guerra-Assunção JA, Glynn JR, Perdigão J, Viveiros M, Portugal I, Pain A, Martin N, Clark TG. 2014. A robust SNP barcode for typing Mycobacterium tuberculosis complex strains. Nature communications 5:4812.

12.	Gygli SM, Loiseau C, Jugheli L, Adamia N, Trauner A, Reinhard M, Ross A, Borrell S, Aspindzelashvili R, Maghradze N. 2021. Prisons as ecological drivers of fitness-compensated multidrug-resistant Mycobacterium tuberculosis. Nature medicine 27:1171-1177.

13.	Nguyen L-T, Schmidt HA, Von Haeseler A, Minh BQ. 2015. IQ-TREE: a fast and effective stochastic algorithm for estimating maximum-likelihood phylogenies. Molecular biology and evolution 32:268-274.

14.	Letunic I, Bork P. 2021. Interactive Tree Of Life (iTOL) v5: an online tool for phylogenetic tree display and annotation. Nucleic acids research 49:W293-W296.

15.	Taneja NK, Tyagi JS. 2007. Resazurin reduction assays for screening of anti-tubercular compounds against dormant and actively growing Mycobacterium tuberculosis, Mycobacterium bovis BCG and Mycobacterium smegmatis. Journal of antimicrobial chemotherapy 60:288-293.

16.	Martin A, Camacho M, Portaels F, Palomino JC. 2003. Resazurin microtiter assay plate testing of Mycobacterium tuberculosis susceptibilities to second-line drugs: rapid, simple, and inexpensive method. Antimicrobial agents and chemotherapy 47:3616-3619.

17.	Rajiniraja M. 2024. 18 Screening of Antimycobacterial. Protocols of Actinomycetes: Microbiology to Gene editing:125.
